# Serum proteomics identifies immune pathways and candidate biomarkers of coronavirus infection in wild vampire bats

**DOI:** 10.1101/2022.01.26.477790

**Authors:** Daniel J. Becker, Guang-Sheng Lei, Michael G. Janech, Alison M. Brand, M. Brock Fenton, Nancy B. Simmons, Ryan F. Relich, Benjamin A. Neely

## Abstract

The apparent ability of bats to harbor many virulent viruses without showing disease is likely driven by distinct immune responses that coevolved with mammalian flight and the exceptional longevity of this order. Yet our understanding of the immune mechanisms of viral tolerance is restricted to a small number of bat–virus relationships and remains poor for coronaviruses (CoVs), despite their relevance to human health. Proteomics holds particular promise for illuminating the immune factors involved in bat responses to infection, because it can accommodate especially low sample volumes (e.g., sera) and thus can be applied to both large and small bat species as well as in longitudinal studies where lethal sampling is necessarily limited. Further, as the serum proteome includes proteins secreted from not only blood cells but also proximal organs, it provides a more general characterization of immune proteins. Here, we expand our recent work on the serum proteome of wild vampire bats (*Desmodus rotundus*) to better understand CoV pathogenesis. Across 19 bats sampled in 2019 in northern Belize with available sera, we detected CoVs in oral or rectal swabs from four individuals (21.1% positivity). Phylogenetic analyses identified all vampire bat sequences as novel α-CoVs most closely related to known human CoVs. Across 586 identified serum proteins, we found no strong differences in protein composition nor abundance between uninfected and infected bats. However, receiver operating characteristic curve analyses identified seven to 32 candidate biomarkers of CoV infection, including AHSG, C4A, F12, GPI, DSG2, GSTO1, and RNH1. Enrichment analyses using these protein classifiers identified downregulation of complement, regulation of proteolysis, immune effector processes, and humoral immunity in CoV-infected bats alongside upregulation of neutrophil immunity, overall granulocyte activation, myeloid cell responses, and glutathione processes. Such results denote a mostly cellular immune response of vampire bats to CoV infection and identify putative biomarkers that could provide new insights into CoV pathogenesis in wild and experimental populations. More broadly, applying a similar proteomic approach across diverse bat species and to distinct life history stages in target species could improve our understanding of the immune mechanisms by which wild bats tolerate viruses.

## Introduction

Bats (Order: Chiroptera) are one of the most speciose mammalian clades (Gunnell and Simmons, 2012; Simmons and Cirranello, 2019), and this diversity is matched by an equally high richness of viruses, many of which are zoonotic (Olival et al., 2017; Mollentze and Streicker, 2020). Bat species are the confirmed reservoir hosts for Hendra and Nipah viruses, lyssaviruses, Marburg virus, and SARS-like coronaviruses (CoVs) (Li et al., 2005; Banyard et al., 2011; Halpin et al., 2011; Amman et al., 2015). Yet with few exceptions (e.g., lyssaviruses), these pathogens seem to not cause disease in their bat hosts (Williamson et al., 2000; Watanabe et al., 2010; Munster et al., 2016). This tolerance of otherwise virulent infections is likely driven by distinct aspects of bat immunity that evolved alongside their unique ability of powered flight and their very long lifespans (Wilkinson and South, 2002; Zhang et al., 2013; Irving et al., 2021). These include but are not limited to constitutive expression of interferons (IFNs) and IFN-stimulated genes (ISGs), robust complement proteins, and a dampened inflammatory response (Zhou et al., 2016; Ahn et al., 2019; Becker et al., 2019; Banerjee et al., 2020; Jebb et al., 2020; Bondet et al., 2021).

Relationships between bats and CoVs specifically have been of particular interest for zoonotic risk assessment (Fischhoff et al., 2021; Becker et al., 2022). CoVs are RNA viruses found across mammals and birds, with at least seven known human viruses: two and five in the genera *Alphacoronavirus* and *Betacoronavirus* (Anthony et al., 2017b; Ye et al., 2020). A novel α-CoV with canine origins was recently detected in humans, but its role in zoonotic disease remains unclear (Vlasova et al., 2021). CoVs are highly diverse in bats, which are the likely ancestral hosts of α- and β-CoVs (Woo et al., 2006, 2012; Ruiz-Aravena et al., 2021). The evolutionary origins of highly pathogenic CoVs (i.e., SARS-CoV, MERS-CoV, SARS-CoV-2) have also been ascribed to bats, but spillover has typically involved intermediate hosts rather than direct bat-to-human transmission (Li et al., 2005; Anthony et al., 2017a; Boni et al., 2020).

Our understanding of bat–CoV interactions and the bat immune response to infection remains limited and has stemmed mostly from experimental studies of a few select species, including *Rousettus leschenaulti, R. aegyptiacus, Artibeus jamaicensis*, and *Eptesicus fuscus* (Watanabe et al., 2010; Munster et al., 2016; van Doremalen et al., 2018; Hall et al., 2021; Ruiz-Aravena et al., 2021). *In vivo* studies have typically found short-term CoV replication and shedding without substantial weight loss or pathology, supporting viral tolerance, although some species seem to entirely resist infection. Tolerance appears driven by innate immune processes, such as increased expression of ISGs, with little adaptive immune response. *In vitro* studies have further supported bat receptor affinity for CoVs (i.e., susceptibility) but little host inflammatory response in bats (Watanabe et al., 2010; Munster et al., 2016; Ahn et al., 2019; Lau et al., 2020).

Despite the clear insights afforded by experiments, model bat systems are heavily limited by logistical constraints (e.g., necessity for specialized facilities, colony maintenance) and have been mostly focused on frugivorous bats, which are relatively easy to keep in captivity compared to other dietary guilds (Banerjee et al., 2019; Wang et al., 2021). Transcriptomics of key tissues (e.g., spleen) has helped advance the field by identifying immune responses to infection in a wider array of species (Papenfuss et al., 2012; Davy et al., 2018; Ren et al., 2020). However, such approaches are restrictive when lethal bat sampling is limited, such as when working with threatened species, in protected habitats, or in longitudinal studies to assess how these viruses persist in bat populations. Blood transcriptomes overcome such challenges to a degree and are increasingly feasible (Huang et al., 2016, 2019), yet such assays are only informative about the immune response in the blood itself. Proteomics, on the other hand, provides a unique and more nuanced perspective into the immune system, because the blood proteome includes proteins secreted from not only blood cells but also proximal organs such as the liver (Uhlén et al., 2019).

Proteomics holds special promise for illuminating the innate and adaptive immune factors involved in bat responses to infection, because it can accomodate the small bat blood volumes typical of field studies (e.g., <10 μL). Recently, we surveyed the serum proteome of wild vampire bats (*Desmodus rotundus*) (Neely et al., 2020). Owing to its diet of mainly mammal blood, this species is the primary reservoir host of rabies lyssavirus in Latin America, and alpha- and betacoronaviruses have also been identified in this species (Brandão et al., 2008; Schneider et al., 2009; Bergner et al., 2020; Alves et al., 2021). Using only 2 μL sera from these small (25– 40 gram) bats, we identified 361 proteins across five orders of magnitude, including antiviral and antibacterial components, 20S proteasome complex, and redox activity proteins. Mass spectrometry–based proteomics can thus facilitate the relative quantification of classical immunological proteins while also providing insight into proteins yet to be fully recognized for their importance in resolving viral infection (Neely et al., 2014, 2018; Geyer et al., 2017).

Here, we expand our recent work on serum proteomics in vampire bats in the context of CoV infection. First, by profiling the serum proteome of the same host species in an additional year of study, we provide a more general and comprehensive characterization of the wild bat immune phenotype. Second, owing to changes in regulations for importing vampire bat samples into the United States, we also assess the impact of heat inactivation, a common method of inactivating bat sera (Schulz et al., 2020), on the proteome. Lastly, and most importantly, we assess differences in the serum proteomes of wild bats with and without acute CoV infection. We aimed to identify up- and downregulated immune responses of wild bats to CoV infection, with particular interest in comparisons with results from experimental infections. We also aimed to guide the discovery of candidate serum biomarkers of viral infection. By taking an agnostic approach via discovery proteomics (Aebersold and Mann, 2016), such biomarkers could provide new and mechanistic insight into CoV pathogenesis in wild bats (Heck and Neely, 2020).

## Materials and Methods

### Vampire bat sampling

As part of an ongoing longitudinal study (Becker et al., 2020b), we sampled vampire bats in April 2019 in the Lamanai Archeological Reserve, northern Belize. This same population was sampled in 2015 for our earlier proteomic analysis (Neely et al., 2020). For the 19 individuals included in this study, we used a harp trap and mist nets to capture bats upon emergence from a tree roost. All individuals were issued a unique incoloy wing band (3.5 mm, Porzana Inc) and identified by sex, age, and reproductive status. For sera, we collected blood by lancing the propatagial vein with 23-gauge needles followed by collection with heparinized capillary tubes. Blood was stored in serum separator tubes (BD Microtainer) for 10−20 minutes before centrifugation. Following recent CDC guidelines, all sera were inactivated for importation to the United States by heating at 56 °C for one hour. We also collected saliva and rectal samples using sterile miniature rayon swabs (1.98 mm; Puritan) stored in DNA/RNA Shield (Zymo Inc). Samples were held at −80 °C using a cryoshipper (LabRepCo) prior to long-term storage. Bleeding was stopped with styptic gel, and all bats were released at their capture location. Field protocols followed guidelines for safe and humane handling of bats from the American Society of Mammalogists (Sikes and Gannon, 2011) and were approved by the Institutional Animal Care and Use Committee of the American Museum of Natural History (AMNHIACUC-20190129). Sampling was further authorized by the Belize Forest Department via permit FD/WL/1/19(09). Serum specimens used for proteomic analysis were approved by the National Institute of Standards and Technology Animal Care and Use Coordinator under approval MML-AR19-0018.

### CoV screening and phylogenetics

As part of a larger viral surveillance project, we extracted and purified RNA from oral and rectal swabs using the *Quick*-RNA Viral Kit (Zymo Research). With exception of one bat, RNA from both swab types was available for all sera. We used a semi-nested PCR targeting the RNA-dependent RNA polymerase gene (RdRp) of alpha- and betacoronaviruses, following previous protocols (Monchatre-Leroy et al., 2017). Amplicons were submitted to GENEWIZ for sequencing. Resulting sequences were aligned using Geneious (Biomatters; (Kearse et al., 2012), followed by analysis using NCBI BLAST (Altschul et al., 1990). PhyML 3.0 was used to build a maximum-likelihood phylogeny of these and additional CoV sequences (Guindon et al., 2010).

To assess possible risk factors for CoV infection (in oral or rectal swab samples), we fit univariate logistic regression models with bat age, sex, and reproductive status as separate predictors. Because the small sample sizes used here could bias our estimates of odds ratios, we used the *logistf* package in R to implement Firth’s bias reduction (Heinze and Schemper, 2002).

### Proteome profiling

In addition to profiling serum from the 19 bats described above, we also performed another proteomic experiment to evaluate effects of heat inactivation with previously analyzed samples collected in 2015 (Neely et al., 2020). From four non-inactivated sera, we submitted an aliquot of each sample to the heat inactivation process used for our 2019 samples (56 °C for one hour). We then processed the paired non-inactivated and heat-inactivated sera and the 19 sera in two batches using the S-Trap method for digestion with the S-Trap micro column (ProtiFi; ≤ 100 μg binding capacity). Full details are provided in the Supplemental Information. Briefly, we used 2 μL (approximately 100 μg protein) of each serum sample for digestion, and a pooled bat serum was digested across the two batches. Proteins were reduced with DL-Dithiothreitol (DTT) and alkylation with 2-chloroacetamide (CAA). Digestion was performed with trypsin at an approximate 1:30 mass ratio, followed by incubation at 47 °C for one hour. These resulting peptide mixtures were then reduced to dryness in a vacuum centrifuge at low heat before long-term storage at −80 °C. Before analysis, samples were then reconstituted with 100 μL 0.1% formic acid (volume fraction) and vortexed, followed by centrifugation at 10000 x *g*_n_ for 10 minutes at 4 °C. The sample peptide concentrations were determined via the Pierce quantitative colorimetric peptide assay with a Molecular Devices SpectraMax 340PC384 microplate reader.

We used the same LC-MS/MS method earlier applied for vampire bat serum proteomics (Neely et al., 2020). Using the original sample randomization gave a randomized sample order, and injection volumes were determined for 0.5 μg loading (0.21–0.44 μL sample). The run order and data key are provided in Table S1. Peptide mixtures were analyzed using an UltiMate 3000 Nano LC coupled to a Fusion Lumos Orbitrap mass spectrometer (ThermoFisher Scientific). A trap elute setup was used with a PepMap 100 C18 trap column (ThermoFisher Scientific) followed by separation on an Acclaim PepMap RSLC 2 μm C18 column (ThermoFisher Scientific) at 40 °C. Following 10 minutes of trapping, peptides were eluted along a 60 minute two-step gradient of 5–30% mobile phase B (80% acetonitrile volume fraction, 0.08% formic acid volume fraction) over 50 minutes, followed by a ramp to 45% mobile phase B over 10 minutes, ramped to 95% mobile phase B over 5 minutes, and held at 95% mobile phase B for 5 minutes, all at a flow rate of 300 nL per minute. The data-independent acquisition (DIA) settings are briefly described here. The full scan resolution using the orbitrap was set at 120000, the mass range was 399 to 1200 *m/z* (corresponding to the DIA windows used), with 40 DIA windows that were 21 *m/z* wide, with 1 *m/z* overlap on each side covering the range of 399 to 1200 *m/z*. Each DIA window used higher-energy collisional dissociation at a normalized collision energy of 32 with quadrupole isolation width at 21 *m/z*. The fragment scan resolution using the orbitrap was set at 30000, and the scan range was specified as 200 to 2000 *m/z*. Full details of the LC-MS/MS settings are in Supplemental Material. The method file (85min_DIA_40×21mz.meth) and mass spectrometry proteomics data have been deposited to the ProteomeXchange Consortium via the PRIDE (Perez-Riverol et al., 2022) partner repository with the dataset identifier PXD031075.

In our earlier analysis of vampire bat serum proteomes (Neely et al., 2020), we used Spectronaut software to analyze our DIA data. Here, we instead used the DIA-NN software suite, which uses deep neural networks for the processing of DIA proteomic experiments (Demichev et al., 2020). To search the bat samples, we used the NCBI RefSeq *Desmodus rotundus* Release 100 GCF_002940915.1_ASM294091v2 FASTA (29,845 sequences). Our full DIA-NN settings are provided in the Supplemental Material, and search settings and the generated spectral library (*.speclib) are included in the PRIDE submission (PXD031075). Briefly, 0.01 precursor false discovery was used, search parameters were chosen based on DIA settings, trypsin (cut at K*,R* but excluded cuts at *P) was selected, and fixed modification of cysteine carbamidomethylation. Since DIA-NN was made to handle protein inference and grouping assuming a UniProtKB-formatted FASTA file, and the NCBI RefSeq *Desmodus rotundus* Release 100 was used (RefSeq format), settings were chosen such that DIA-NN effectively ignored protein grouping, which can be performed on the backend following ontology mapping.

To additionally search for CoV proteins (Neely et al., 2020), we performed a secondary search using the same settings and the addition of a Coronaviridae FASTA (117709 sequences) retrieved from UniProtKB (2021_03 release) using taxon identifier 1118 with all SwissProt and TrEMBL entries. Search settings are included in the PRIDE submission (PXD031075).

Our identified bat proteins were then mapped to human orthologs using BLAST+ (Camacho et al., 2009) and a series of python scripts described previously (Neely et al., 2020) to facilitate downstream analysis using human-centric databases (see Supplemental Material for full details). In those cases where human orthologs do not exist, such as mannose-binding protein A (MBL1), we used *ad hoc* ortholog identifiers. Specifically, eight identified vampire bat proteins are not found in humans (APOR, Bpifb9a, HBE2, ICA, LGB1, MBL1, Patr-A, and REG1), and we thus used UniProt identifiers from chimpanzee, cow, horse, mouse, and pig (Table S2).

### Proteomic data analyses

The final data matrix of relative protein abundance for all identified proteins was stratified into two datasets for differential analysis: (*i*) the four 2015 samples analyzed before and after heat inactivation (Table S3) and (*ii*) the 19 samples collected in 2019 analyzed for CoV infection (Table S4). Our analysis also included a pooled serum sample as a quality control between the two digestion batches (Table S2), and digestion was evaluated by the number of peptide spectral matches. For formal statistical analyses, missing abundance values were imputed as half the minimum observed intensity of each given protein; however, for summary statistics (e.g., means, log_2_-fold change [LFC]), missing values were excluded (Lazar et al., 2016; Arioli et al., 2021).

For the inactivation analyses, log_2_-transformed protein abundance ratios were used for each paired sample. These ratios were used in a moderated *t*-test with the *limma* package in R to evaluate protein changes within sera samples before and after heat treatment (Ritchie et al., 2015), followed by Benjamini–Hochberg (BH) correction (Benjamini and Hochberg, 1995).

For the CoV infection analyses, we first reduced dimensionality of our protein dataset using a principal components analysis (PCA) of all identified proteins, with abundances scaled and centered to have unit variance. We then used a permutation multivariate analysis of variance (PERMANOVA) with the *vegan* package to test for differences in protein composition between uninfected and infected bats (Dixon, 2003). Next, we used a two-sided Wilcoxon rank sum test in MATLAB to detect differentially abundant proteins between uninfected and infected bats. We again used the BH correction to adjust for the inflated false discovery rate. We also calculated LFC as the difference of mean log_2_-transformed counts between uninfected and infected bats. To next identify candidate biomarkers of CoV infection, we used receiver operating characteristic (ROC) curve analysis. We used a modified function (https://github.com/dnafinder/roc) in MATLAB to generate the area under the ROC curve (AuROC) as a measure of classifier performance with 95% confidence intervals, which we calculated with standard error, α = 0.05, and a putative optimum threshold closest to 100% sensitivity and specificity (Hanley and McNeil, 1982; Pepe, 2003). We considered proteins with AuROC ≥ 0.9 to be strict classifiers of CoV positivity, whereas proteins with AuROC ≥ 0.8 but less than 0.9 were considered less conservative; all other proteins were considered to be poor classifiers (Mallick et al., 2007). Variation in the abundance of strict classifiers by CoV infection status was visualized using boxplots. We also visualized the matrix of all candidate serum biomarkers with the *pheatmap* package, using log_2_-transformed protein abundances (scaled and centered around zero) and Ward’s hierarchical clustering method (Murtagh and Legendre, 2014; Kolde and Kolde, 2015).

Lastly, we interrogated up- and down-regulated responses to CoV infection using gene ontology (GO) analysis. First, we programmatically accessed GO terms for all identified proteins using their associated UniProt identifiers and the *UniprotR* package (Soudy et al., 2020). Next, we used the *gprofiler2* package as an interface to the g:Profiler tool g:GOSt for functional enrichment tests (Raudvere et al., 2019; Kolberg et al., 2020). Enrichment was performed for all candidate protein biomarkers based on AuROC, with up- and downregulated proteins determined using log_2_-fold change. We ranked our proteins by AuROC to conduct incremental enrichment testing, with the resulting *p-*values adjusted by the Set Counts and Sizes (SCS) correction. We restricted our data sources to GO biological processes, the Kyoto Encyclopedia of Genes and Genomes (KEGG), and WikiPathways (WP). We ran the enrichment tests for both our strict and less-conservative protein classifiers. We note that the eight bat proteins lacking human orthologs all had AuROC < 0.8 and therefore did not require any manual GO and pathway mapping.

## Results

### Bat demographics and CoV positivity

Our sample of 19 vampire bats included for proteomic analyses consisted predominantly of females (84 %) and adults (79 %). One male was reproductively active, whereas four females were lactating (*n* = 3) or pregnant (*n* = 1). Four of the 19 sera samples had paired oral or rectal swabs test positive through PCR for CoVs (21.1 %); sequences are available in GenBank under accession numbers OM240577−80. Phylogenetic analyses of the four sequenced amplicons confirmed all positives in the genus *Alphacoronavirus* (Fig. 1A). All four vampire bat sequences are novel α-CoVs and displayed the most genetic similarity (94.6−97.3 %) to human CoVs (HCoVs; HCoV-NL63 and HCoV-229E) rather than known bat α-CoVs, including those previously found in other vampire bat colonies and other Neotropical bat species more broadly (Asano et al., 2016; Bittar et al., 2020). Univariate logistic regressions did not find significant (unadjusted) effects of any bat demographics on CoV positivity. Males were no more likely than females to be infected (OR = 2.31, *p* = 0.49), although non-reproductive individuals (OR = 6.16, *p* = 0.17) and subadults (OR = 5.40, *p* = 0.13) were weakly more likely to be positive (Fig. 1B).

**Figure 1.**
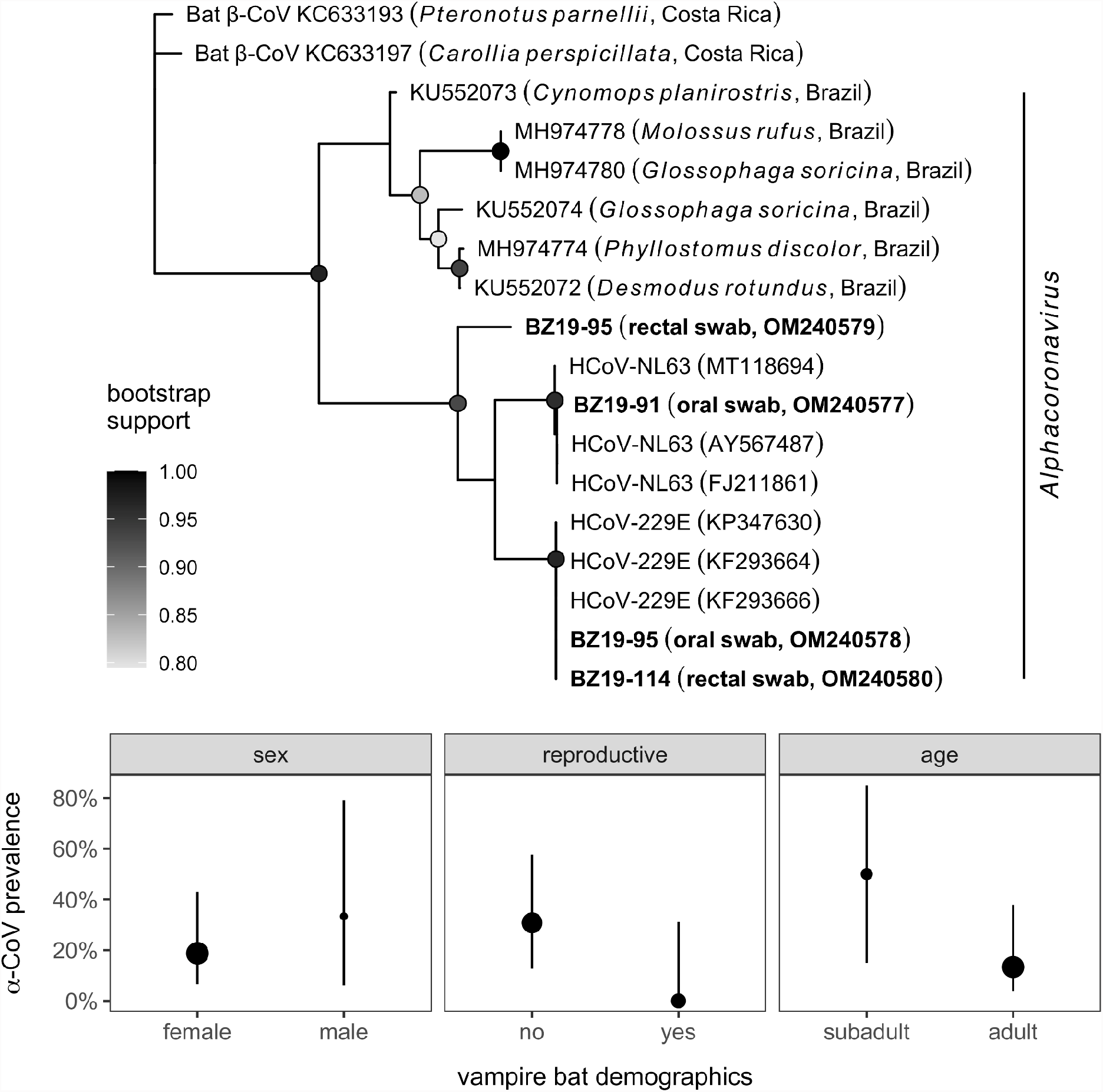
Phylogeny and distribution of CoV infections in Belize vampire bats. (A) Sequences from PCR-positive oral or rectal swabs (*n* = 4) were aligned with genetically similar (NCBI BLAST) and representative CoV sequences from other Neotropical bat species. Nodes display bootstrap support from maximum likelihood estimation; only values greater than 80% are shown. (B) Prevalence of alphacoronavirus infection in bats with paired sera samples alongside 95% confidence intervals (Wilson’s method) for key bat demographic variables. Point estimates of infection prevalence (oral and rectal swabs pooled) are scaled by sample size per covariate.

### Serum proteome characterization

Bottom-up proteomics using DIA identified 586 proteins in the 19 vampire bat sera samples, with relative quantification covering 5.6 orders of magnitude (Fig. 2A; Table S4). The overall number of identified proteins in our former analysis (i.e., 361 proteins) (Neely et al., 2020) and the current dataset was within the same order of magnitude and had a similar dynamic range (approximately 10^3^−10^8^ in our prior analysis versus 10^4^−10^9^ in the current analysis). There was also high overlap in identified proteins between the datasets (91 % of the original 361 proteins were included in our analysis here; Fig. S1) and similar protein ranks (Table S5). Although the prior and current study have low sample sizes (*n* = 17 and 19, respectively) and were sampled across different years, the similarity in protein abundance, composition, and ranks suggest that these proteomic patterns (e.g., guanylate-binding proteins, circulating 20S proteasome, and hyaluronidase-1) are not the result of sampling bias and are instead likely a consistent vampire bat phenotype, with improved protein identifications differences driven by technical advances.

**Figure 2.**
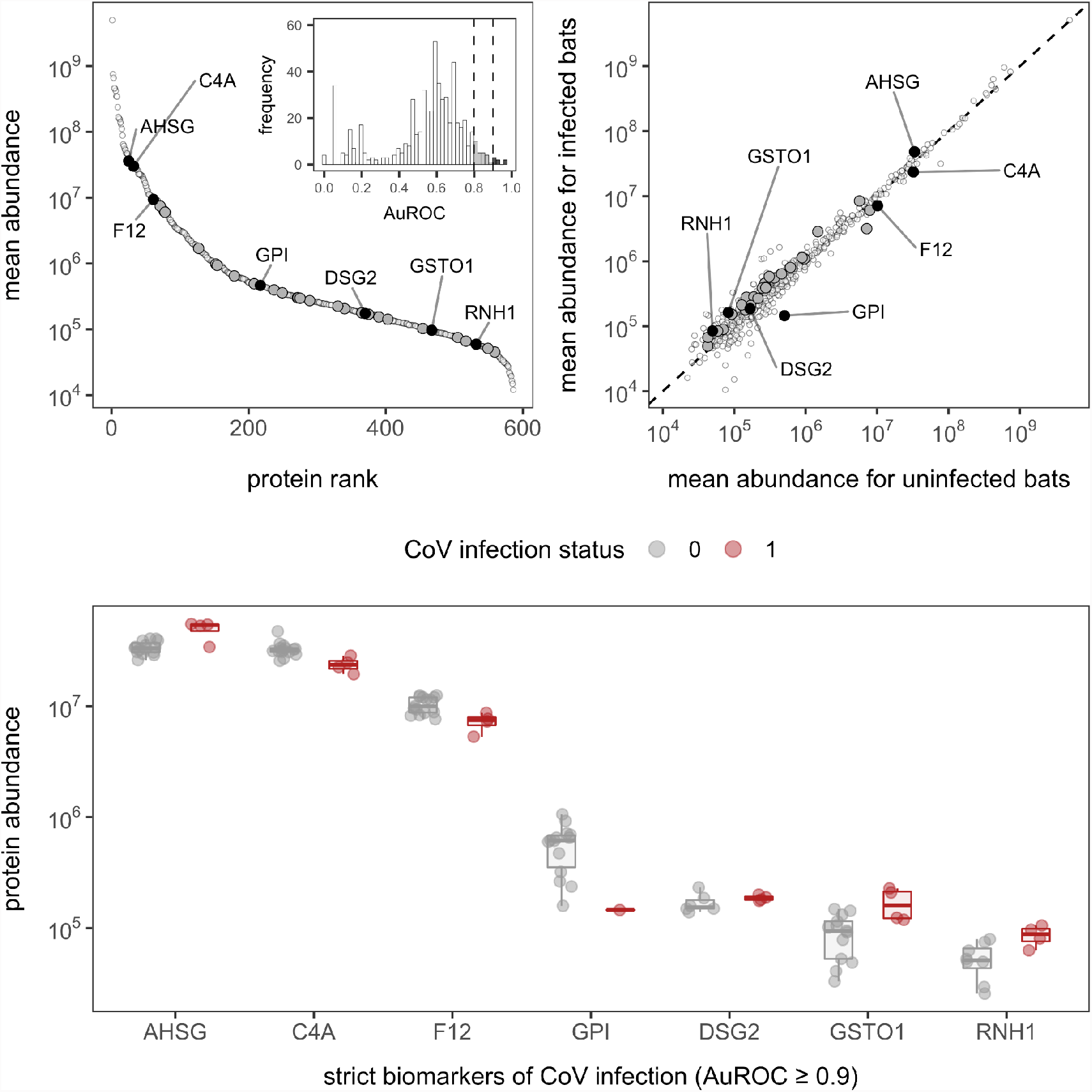
Serum protein abundance and biomarkers of CoV infection in vampire bats. (A) Mean abundance of the 586 proteins was plotted with corresponding rank to illustrate the dynamic range of the serum proteome. The inset displays the distribution of AuROC values for protein biomarkers of CoV infection alongside cutoffs of 0.8 and 0.9 for strict and less-conservative biomarkers. (B) Comparison of mean protein abundance for uninfected and infected bats across all proteins; the dashed line shows the 1:1 reference. For both plots, less-conservative and strict biomarkers are shown in dark gray and black, with the latter labeled with gene symbols (AHSG, alpha-2-HS-glycoprotein; C4A, complement C4A; F12, coagulation factor XII; GPI, glucose-6-phosphate isomerase; DSG2, desmoglein-2; GSTO1, glutathione S-transferase omega-1; RNH1, ribonuclease inhibitor). Missing values were excluded prior to determining mean abundances. (C) Protein abundance between uninfected and infected bats for strict biomarkers of CoV positivity; boxplots are overlaid by raw data jittered to reduce overlap.

### Effects of heat inactivation

One key difference between analyses here and our prior proteomic study of this vampire bat population is unknown technical artifacts from heat inactivation. To assess these possible effects, we compared proteomes before and after treatment of four serum samples used in our prior study (Neely et al., 2020). Using a moderated *t*-test of the four paired sera samples, 34 proteins showed significant changes after heat inactivation (unadjusted *p* < 0.05), but no differences remained after BH adjustment (even using a liberal adjusted *p* < 0.3). Although we found no statistically significant changes in protein abundance with heat inactivation, we observed a mean 28% absolute change across the proteome, with a maximum mean 500% absolute change. Most proteins (52.6%) changed less than 17% in response to heat inactivation (Fig. S2; Table S3).

### Mining for CoV proteins

Given prior proteomic identification of putative viral proteins in undepleted serum, including CoVs (Neely et al., 2020), we broadened our search space to include any CoV proteins. As observing non-host proteins is a rare event, we used additional stringent criteria to verify any initial CoV peptide spectral matches (see Supplemental Material). Of the 749 CoV peptide spectral matches, none passed these more stringent criteria (Table S6). Thus we cannot firmly say that viral proteins were identified in this set of undepleted sera, regardless of CoV status.

### Proteomic differences with CoV infection

To assess differences in the serum proteome between CoV-infected and uninfected bats, we first used multivariate tests. Across the 586 identified proteins, the first two PCs explained 25.46 % of the variance in serum proteomes (Fig. S3). A PERMANOVA found no difference in proteome composition by viral infection status (*F*_*1,17*_ = 0.99, *R*^*2*^ = 0.05, *p* = 0.46), although variation was greater in infected bats. Using Wilcoxon rank sum tests, we initially identified 22 proteins with significantly different abundance in CoV-infected bats (unadjusted *p* < 0.05), but no differences likewise remained after BH adjustment (even using a liberal adjusted *p* < 0.3; Fig. 2B).

In contrast to multivariate and differential abundance tests, ROC curve analyses identified 32 candidate protein biomarkers of CoV infection using strict (*n* = 7, AuROC ≥ 0.9) and less-conservative (*n* = 25, 0.9 > AuROC ≥ 0.8) classifier cutoffs (Fig. 2). Considering the strict classifier cutoff, four *Desmodus* proteins were positive predictors and were weakly elevated in CoV-infected bats: RNH1 (ribonuclease inhibitor; AuROC = 0.97, LFC = 0.77), AHSG (alpha-2-HS-glycoprotein; AuROC = 0.91, LFC = 0.52), DSG2 (desmoglein-2; AuROC = 0.90, LFC = 0.16), and GSTO1 (glutathione S-transferase omega-1; AuROC = 0.90, LFC = 0.98). Conversely, three proteins were instead negative predictors and were reduced in CoV-infected bats: C4A (complement C4A; AuROC = 0.97, LFC = −0.45), F12 (coagulation factor XII; AuROC = 0.93, LFC = −0.51), and GPI (glucose-6-phosphate isomerase; AuROC = 0.03, LFC = −1.79; Fig. 2C). The total 32 candidate biomarkers provided clear discriminatory power in differentiating the phenotypes of uninfected and infected vampire bats (Fig. 3).

**Figure 3.**
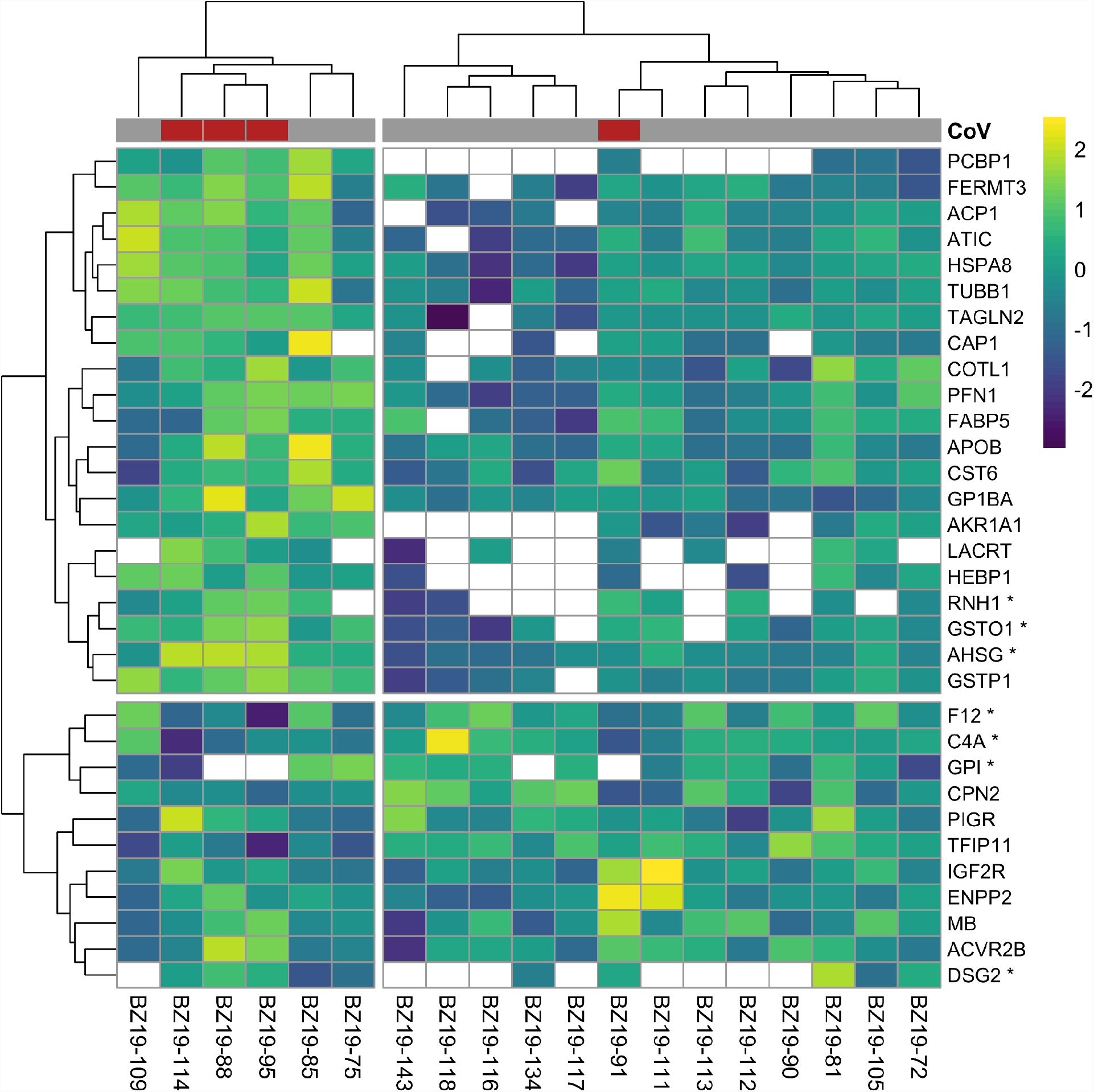
Heatmap of log_2_-transformed abundance for all candidate serum biomarkers of CoV infection (*n* = 32), scaled to a mean of zero. Rows display individual bats, while columns display proteins as gene symbols; those with AuROC ≥ 0.9 are marked with an asterix. CoV infection status is shown at the top of the heatmap. Clustering used Ward’s hierarchical method (Murtagh and Legendre, 2014; Kolde and Kolde, 2015). Missing abundance values are shown as blank.

We lastly interrogated up- and down-regulated responses to CoV infection using GO terms. Across the serum proteome (*n* = 586), top biological processes (≥ 30 proteins; Fig. S4) included neutrophil degranulation (18.3 %), platelet degranulation (8.5 %), post-translational protein modification (8.5 %), innate immune response (7.3 %), cellular protein metabolic process (6.5 %), blood coagulation (5.8%), cell adhesion (5.5 %), signal transduction (5.5 %), viral processes (5.5 %), negative regulation of apoptotic process (5.3 %), inflammatory response (5.1 %), and regulation of complement activation (5.1 %). Enrichment analyses identified multiple functional protein differences between uninfected and infected bats after SCS correction (Fig. 4). When considering only strict protein biomarkers (AuROC ≥ 0.9), CoV-infected bats displayed downregulation of the complement system and regulation of proteolysis. When also considering less-conservative biomarkers (AuROC ≥ 0.8), infected bats also had down-regulation of immune effector processes and humoral immunity. Enrichment analysis of all biomarkers also identified up-regulated processes including neutrophil-mediated immunity, overall granulocyte activation, myeloid cell responses, and glutathione processes, denoting a largely cellular immune response.

**Figure 4.**
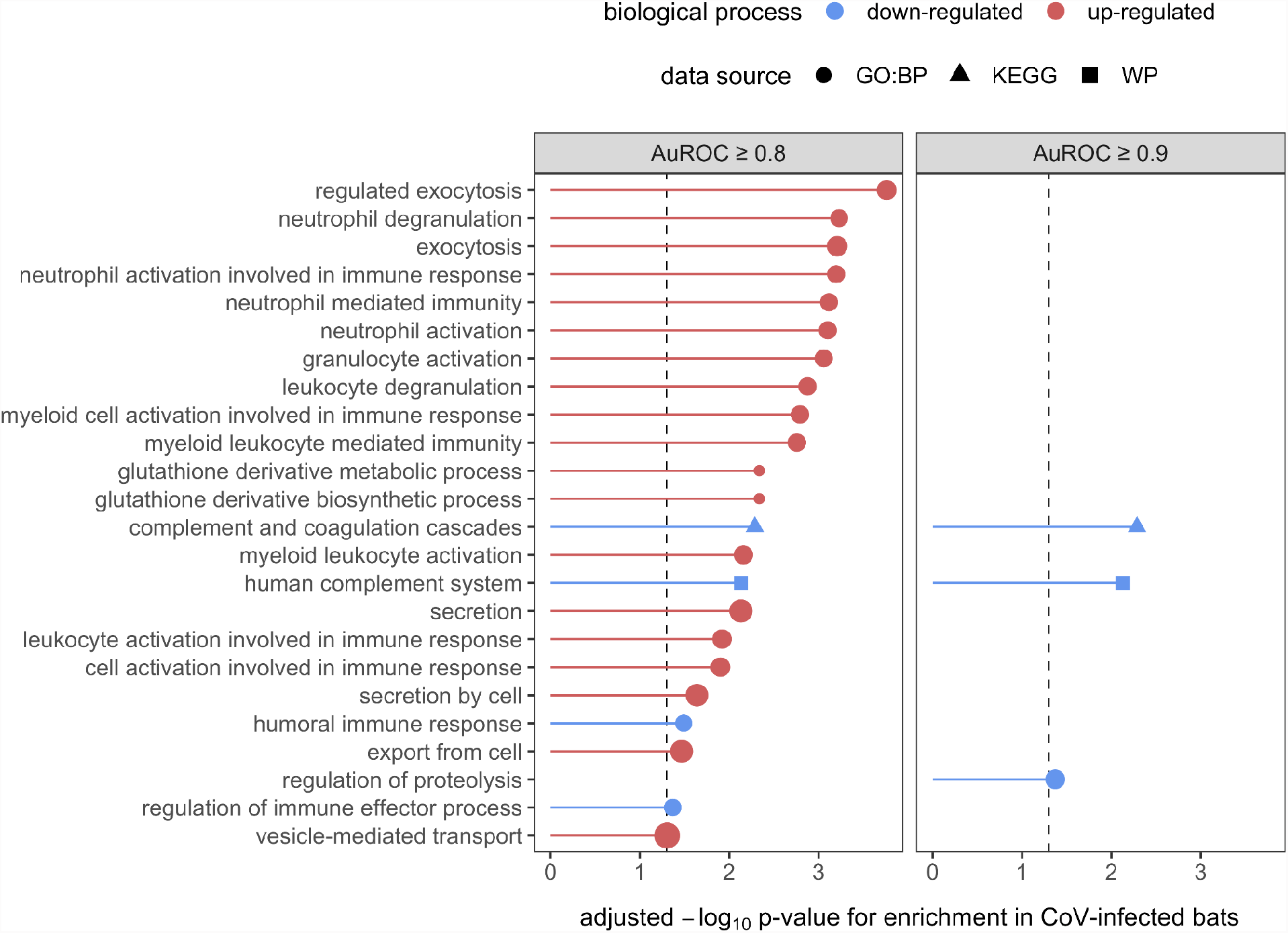
Enrichment analyses of the 32 candidate biomarkers of CoV infection, stratified by less-conservative (AuROC ≥ 0.8) and strict (AuROC ≥ 0.9) classifiers. Biological processes with significant enrichment in CoV-infected bats after SCS correction are displayed, with up- and downregulated processes shown in red and blue, respectively. Processes are labeled by source.

## Discussion

Despite an increasing interest in bat–virus interactions, especially for CoVs given their human health relevance (Anthony et al., 2017b; Becker et al., 2022), we still have limited insights into the immune mechanisms involved in infection of bats (Ruiz-Aravena et al., 2021). Here, we used serum proteomics to broadly profile the immune phenotype of wild vampire bats in the presence of relatively common CoV infections. Novel α-CoVs detected in these bats had little association with serum protein composition nor abundance, although ROC curve analyses identified 7–32 candidate biomarkers of CoV infection, including AHSG, C4A, F12, GPI, DSG2, GSTO1, and RNH1. Enrichment analyses using these protein classifiers identified strong downregulation of complement, regulation of proteolysis, immune effector processes, and humoral immunity in CoV-infected bats alongside an upregulation of neutrophil immunity, overall granulocyte activation, myeloid cell responses, and glutathione processes. Such results denote a mostly cellular immune response of vampire bats to CoVs and further identify putative biomarkers that could provide new insights into CoV pathogenesis in both wild and experimental populations.

Much bat immunology work to date has understandably focused on model bat systems under captive conditions (Banerjee et al., 2020; Irving et al., 2021). However, identifying the immune correlates of infection is especially important for wild populations, where susceptibility and tolerance to infection, alongside other immunological processes, can vary based on habitat quality and host life history (e.g., reproduction) (Becker et al., 2020a). Such efforts could provide a mechanistic basis for establishing when and where pathogen pressure from bats is greatest and thus help predict viral spillover (Plowright et al., 2016; Becker et al., 2021). Unlike some other global profiling techniques, proteomics has the benefit of leveraging the small blood volumes that can be obtained non-lethally from most small bats (e.g., members of the Yangochiroptera, including over 1000 species) and thus is especially amenable to the long-term, mark–recapture studies required to study bat virus dynamics (Plowright et al., 2019; Heck and Neely, 2020). To facilitate this work, we here built on our prior proteomic characterization of *Desmodus rotundus* (Neely et al., 2020). We identified a core serum protein phenotype for this species from multiple years of sampling that could serve as a reference for long-term proteomic studies, although technical advances (e.g., DIA-NN) likely contributed to an expanded protein repertoire in the current study. We also assessed the impacts of heat treatment, a common inactivation method for sera, given recent shifts in United States importation regulations. Artifacts from heat inactivation were not sufficiently conserved to be statistically significant, and most serum proteins had small changes in abundance before and after this treatment. Yet given the extent of such changes, we suggest original samples should typically not be analyzed with heated samples for comparative purposes. In contexts where sera inactivation is required, however, heat treatment across samples should not bias characterization of bat serum proteomes. Such optimizations could next be applied across longitudinal timepoints to more broadly study bat–virus interactions in the wild.

We here focused this initial study on CoVs, which have been previously characterized as genetically diverse (α- and β-CoVs) in Neotropical bats, including but not limited to *Desmodus rotundus* (Brandão et al., 2008; Anthony et al., 2013; Corman et al., 2013; Bergner et al., 2020). Our detection of CoVs from a northern Belize colony of vampire bats in swab samples with paired sera (4/19) was higher than other CoV surveys in this species (Anthony et al., 2013; Asano et al., 2016; Wray et al., 2016; Alves et al., 2021), although future studies with larger sample sizes are needed to test if this represents a true geographic difference in virus prevalence. Yet despite many available CoV sequences in GenBank from diverse bat surveys, all CoVs detected here fell within the genus *Alphacoronavirus* but outside of known bat α-CoV clades. Instead, these viruses were more closely related to human α-CoVs, specifically HCoV-NL63 and HCoV-229E, suggesting greater genetic diversity of CoVs within vampire bats than previously recognized. Similarly, such results further suggest the possibility of high zoonotic potential (or spillback) for vampire bat CoVs, which likely varies based on geography given the high degree of genetic differentiation across the broad distribution of *Desmodus rotundus* (Martins et al., 2009; Streicker et al., 2016; Bergner et al., 2021). Such findings should be confirmed with larger sample sizes and characterization, including *in vitro* assessments and attempts at virus isolation.

Despite identifying CoV infection in a small number of bat samples, we were unable to detect CoV proteins in the serum proteomes. Previously, we detected two CoV peptides in sera from this population, but these were likely at the edge of detection limits (Neely et al., 2020). As such detection limits are susceptible to technical artifacts, including but not limited to sample handling, protein processing, or instrument performance, heat inactivation could have affected our ability to identify similar peptides in these bat samples, especially for bats positive for CoVs by PCR. Additionally, our ability to detect viral proteins may have been further restricted by ongoing limitations in applying proteomics to wild species. In humans, over 3000 serum proteins can be detected by mass spectrometry after depletion of the most abundant proteins (Uhlén et al., 2019). However, using antibody-based depletion techniques is not an effective strategy in non-human mammals (Neely et al., 2014), such that undepleted serum proteomics in bats will be limited to the top 300–600 proteins, with false negatives for low abundance proteins such as those of viruses (Anderson and Anderson, 2002). Alternatively, lack of detection of CoV proteins in sera despite detection of CoV RNA in oral and rectal swabs could indicate tropism, as CoVs have been more readily detected in bat feces and saliva than in blood (Smith et al., 2016).

Using our novel α-CoVs, we then tested for differential composition and abundance of serum proteins between uninfected and infected vampire bats. In both cases, we found negligible overall differences in serum proteomes with CoV infection. However, such null results should be qualified by the challenges posed to differential abundance tests by sample imbalance, given the small number of infected relative to uninfected bats (Yang et al., 2006). To partly address this imbalance, we used ROC curve analyses to identify proteins with strict (AuROC ≥ 0.9; *n* = 7) and less conservative (AuROC ≥ 0.8; *n* = 25) classifier ability for infection (Arthur et al., 2014; Neely et al., 2014). The unbiased query of proteins in relation to infection through proteomics can in turn result in detection of unexpected candidate biomarkers and new insights into CoV pathogenesis in bats. For example, we identified increased ribonuclease inhibitor (RNH1) as a putative biomarker. RNH1 inhibits RNase 1 and blocks extracellular RNA degradation, possibly resulting in increased tumor necrosis factor (TNF)–α activation (Zechendorf et al., 2020). Prior cell line studies of *Eptesicus fuscus* have shown limited production of TNF-α upon stimulation (Banerjee et al., 2017), whereas those of *Pteropus alecto* have suggested an induced TNF-α response (Ahn et al., 2019). Whether greater abundance of pro-inflammatory cytokines such as TNF-α occur with CoV infection in bats would thus be a fruitful area for future work based on these RNH1 differences. Similarly, we identified lower complement C4A (one of two C4 isotypes) as another putative biomarker. In humans, lower complement C4A and C3 can signal elevated autoimmunity (Walport, 2002; Wang and Liu, 2021), and decreased complement C4 and C3 in COVID-19 patients also corresponds to disease severity (Zinellu and Mangoni, 2021). The processes that shape serum complement, namely complement synthesis, activation, and clearance, remain poorly characterized in bats (Becker et al., 2019), but the identification of C4A as a classifier could suggest specific explorations into how complement affects CoV infection.

Other candidate biomarkers also had more direct implications for the antiviral response in bats. AHSG (alpha-2-HS-glycoprotein) is a negative acute phase reactant (Lebreton et al., 1979) and here was a positive predictor of CoV infection. In humans, elevated AHSG is accordingly protective against progression of disease caused by SARS-CoV (Zhu et al., 2011), and decreased inflammation could also contribute to viral tolerance in bats. We also identified poly(rC)-binding protein 1 (PCBP1) as a positive, albeit weaker, predictor of CoV infection (AuROC = 0.87). This RNA-binding protein is upregulated in activated T cells to control effector T cells converting into regulatory T cells and thus stabilizes the innate immune response (Ansa-Addo et al., 2019, 2020); PCBP1 may also prevent virus-related inflammation (Zhou et al., 2012). Whereas human patients with prolonged SARS-CoV-2 infections showed lower PCB1 compared to patients with short-term infections (Yang et al., 2021), bats with CoV infection here had elevated PCBP1 and more generally harbor more PCBP1 than humans (Neely et al., 2020). Despite focusing on two different viral genera, such results suggest both similarities and differences in how bats and humans may respond to CoV infection. More generally, the list of such candidate biomarkers identified here can be used to create accurate, sensitive, quantitative, and bat-specific parallel reaction monitoring mass spectrometry-based protein assays (Neely et al., 2013, 2015). Such assays could facilitate more thorough investigations into bat immune response to CoV infection.

In addition to identifying candidate biomarkers, we also leveraged these proteins to more generally assess broad up- and downregulated biological processes with CoV infection through enrichment analyses. Using all candidate biomarkers, we found that CoV-infected bats displayed downregulation of the complement system, regulation of proteolysis, immune effector processes, and humoral immunity while also showing upregulated neutrophil-mediated immunity, overall granulocyte activation, myeloid cell responses, and glutathione processes. These results in part support findings from limited experimental infections of select bat species, which have shown little humoral response to CoVs (Munster et al., 2016; van Doremalen et al., 2018). Yet while *Rousettus aegyptiacus* challenged with SARS-like CoVs did not show hematological changes following infection (van Doremalen et al., 2018), CoV-infected vampire bats here had largely upregulated cellular immune responses. Similarly, experimental studies have suggested bat tolerance of CoVs to be driven by upregulation of cytokine responses and a downregulated inflammatory response (Munster et al., 2016; Ahn et al., 2019; Lau et al., 2020), but we did not find GO terms related to cytokines or inflammation in our analyses. Such discrepancies could again result from our ability to only detect the top 300–600 proteins without antibody-based depletion, which could cause low-abundance proteins (including but not limited to IFNs) being especially difficult to characterize here (Anderson and Anderson, 2002). Alternatively, these differences could reflect distinct immune responses of bats for α-CoV infection, given that experimental studies to date have focused on β-CoVs. Additionally, these findings could also signal immunological variation within and among bat clades, given that *Desmodus rotundus* and the closely related *Artibeus jamaicensis* may respond differently to CoVs (Munster et al., 2016).

Future proteomic analyses across bat species in the wild could provide a tractable means to broadly characterize host responses to viruses, including but not limited to hypothesized immune mechanisms of tolerance in this order of mammals and to infection with diverse CoVs. By leveraging the benefits of proteomics to quantify hundreds of proteins from the small sera volumes that can be obtained from most bat species (Uhlén et al., 2019; Neely et al., 2020), such analyses could evaluate whether particular immune responses to viruses such as CoVs are conserved across bats (e.g., downregulation of humoral immunity) and which may be a feature of particular bat clades. In particular, further comparative proteomic analyses across Neotropical bats, including both additional members of the Phyllostomidae as well as sister families such as the Mormoopidae (Rojas et al., 2016), would illuminate whether vampire bats have particular immunological relationships with CoVs that may facilitate viral tolerance. As suggested in our work here on *Desmodus rotundus*, such studies could also identify putative biomarkers that may suggest novel mechanisms of pathogenesis and facilitate development of protein-specific assays to improve the resource base for studying the immunology of wild bats and bat–virus dynamics.

## Author contributions

DJB, MBF, and NBS conducted fieldwork; GL and RFR ran CoV diagnostics; MGJ, AMB, and BAN conducted proteomic analyses and bioinformatics; DJB and BAN analyzed data; and DJB and BAN wrote the manuscript with contributions from all coauthors.

## Acknowledgements

For assistance with bat sampling logistics and research permits, we thank Mark Howells, Melissa Ingala, Kelly Speer, Neil Duncan, and the staff of the Lamanai Field Research Center. Identification of certain commercial equipment, instruments, software, or materials does not imply recommendation or endorsement by the National Institute of Standards and Technology, nor does it imply that the products identified are necessarily the best available for the purpose.

## Data availability statement

The raw mass spectrometry proteomics data, associated LC-MS/MS method files, spectral libraries, FASTA, and metadata have been deposited to the ProteomeXchange Consortium via the PRIDE partner repository (Perez-Riverol et al., 2022) with the dataset identifier PXD031075. Vampire bat CoV sequences are available in GenBank under accession numbers OM240577−80.

