## Supplemental Material for "Serum proteomics identifies immune pathways and candidate biomarkers of coronavirus infection in wild vampire bats"

**S1 Full proteomic methods**

**S2 Supplemental tables**

**S3 Supplemental figures**

**S1 Full proteomic methods**

The S-Trap method was used for digestion with the S-Trap micro column (ProtiFi; ≤ 100 µg binding capacity), and the specific method is described in detail below. Based on prior work with other mammalian sera (Neely et al., 2013, 2014), we assumed a protein concentration of approximately 50 µg/µL; this value can vary but assumes at least 50 µg/µL was used for estimating enzyme for the digest. Therefore, 2 µL (approximately 100 µg protein) of each sample was mixed with 48 µL of 50 mmol/L ammonium bicarbonate and 50 µL of 2x lysis buffer [10 % sodium dodecyl sulfate (SDS) volume fraction in 100 mmol/L triethylammonium bicarbonate (TEAB), pH 8.5]. Four non-inactivated sera were used to evaluate heating (since serum samples were heat inactivated prior to shipping, as described in paper). A 2 µL aliquot was removed from each of these four samples and heated at 56 °C for 1 hour, and a companion 2 µL aliquot was removed. These 2 µL heat-inactivated sera and companion non-heated sera were used for digestion steps below. Note that the lysis buffer was added directly to each tube, versus an additional centrifugation step after heating. All samples were reduced with 10 µL of 90 mmol/L DL-Dithiothreitol (DTT; final concentration of 10 mmol/L) at 60 °C for 30 min, then cooled and alkylated with 10 µL of 200 mmol/L 2-chloroacetamide (CAA; final concentration of 20 mmol/L) at room temperature in the dark for 30 min. The sample was acidified with 12 µL of 12% phosphoric acid (volume fraction) bringing the final volumetric ratio to 1:10. Next, 700 µL binding buffer [ProtiFi; 5 % TEAB volume fraction in methanol] was added (approximately 1:7 volumetric ratio)]. Using a vacuum manifold, the sample was washed across the S-Trap with six sequential washes of 400 µL binding buffer. Next, 3 µL of 1 µg/µL trypsin (Pierce) was mixed with 122 µL, 50 mmol/L ammonium bicarbonate, and this 125 µL solution was added to each S-Trap, yielding approximately a 1:30 mass ratio (trypsin:total protein). Samples were incubated at 47 °C for 1 hour, after which they were sequentially washed into 1.5 mL Lo-Bind microcentrifuge tubes (Eppendorf) by centrifugation with the following wash steps: 80 µL 50 mmol/L ammonium bicarbonate, 80 µL 0.2 % formic acid (volume fraction), 80 µL 0.2% formic acid in 50 % acetonitrile (volume fractions) at 1000 x $g_n$, 1000 x $g_n$, and 4000 x $g_n$, respectively at 4 °C for 1 minute. The resulting peptide mixtures were reduced to dryness in a vacuum centrifuge at low heat and stored at –80 °C. Prior to analysis, samples were reconstituted with 100 µL 0.1% formic acid (volume fraction) and briefly vortexed, then centrifuged 10000 x $g_n$ for 10 minutes at 4 °C. The peptide concentration of each sample was determined using the Pierce quantitative colorimetric peptide assay with a Molecular Devices SpectraMax 340PC384 microplate reader.

The same data-independent acquisition (DIA) method previously used for vampire bat serum proteomics (Neely et al., 2020) was used and described in full here. Peptide mixtures in 0.1% formic acid (volume fraction) were analyzed using an UltiMate 3000 Nano LC coupled to a Fusion Lumos Orbitrap mass spectrometer (Thermo Fisher Scientific). Using the original sample randomization yielded a randomized sample order, and injection volumes were determined for 0.5 µg loading (between 0.21 and 0.44 µL sample). The run order and data key are provided in Table S1. Peptide mixtures were loaded onto a PepMap 100 C18 trap column (75 µm id x 2 cm length; Thermo Fisher Scientific) at 3 µL/minutes for 10 minutes with 2% acetonitrile (volume fraction) and 0.05% trifluoroacetic acid (volume fraction) followed by separation on an Acclaim PepMap RSLC 2 µm C18 column (75 µm id x 25 cm length; Thermo Fisher Scientific) at 40 °C. Peptides were separated along a 60 minute two-step gradient of 5% to 30% mobile phase B (80% acetonitrile volume fraction, 0.08% formic acid volume fraction) over 50 minutes followed by a ramp to 45% mobile phase B over 10 minutes and lastly ramped to 95% mobile phase B over 5 minutes, and held at 95% mobile phase B for 5 minutes, all at a flow rate of 300 nL/minute.

The Fusion Lumos was operated in positive polarity with a DIA method constructed using the targetedMS2 module (as opposed to the built-in DIA module). The full scan resolution using the orbitrap was set at 120000, the mass range was 399 to 1200 *m/z* (corresponding to the DIA windows used), 30% RF lens was set, and the full scan ion target value was 4.0e5, allowing a maximum injection time of 20 ms. A default charge of 4 was set under MS Global Settings. As stated, DIA windows were constructed using the targetedMS2 module. Each window used higher-energy collisional dissociation (HCD) at a normalized collision energy of 32 with quadrupole isolation width at 21 *m/z*. The fragment scan resolution using the orbitrap was set at 30 000, the scan range was specified as 200 to 2000 *m/z*, ion target value of 1.0e6 and 60 ms maximum injection time. Data were collected as profile data in both MS1 and MS2, though the authors wish to note that using centroid data is possible for most DIA software. The DIA window scheme was an overlapping static window strategy such that each of the 40 windows were 21 *m/z* wide, with 1 *m/z* overlap on each side covering the range of 399 to 1200 *m/z*. The window centers were specified in the mass list table (with z = 2), such that they were 409.5, 429.5, 449.5, ..., 1189.5. The method file (85min_DIA_40x21mz.meth) and mass spectrometry proteomics data have been deposited to the ProteomeXchange Consortium via the PRIDE (Perez-Riverol et al., 2022) partner repository with the dataset identifier PXD031075.

Previously, Spectronaut software was used to analyze this type of data (Neely et al., 2020), but DIA-NN (Data-Independent Acquisition by Neural Networks) was used for the current analysis (Demichev et al., 2020). The FASTA file used for searching bat samples was the NCBI RefSeq *Desmodus rotundus* Release 100, GCF_002940915.1_ASM294091v2 (29 845 sequences). DIA-NN 1.8 was used for searching and relative quantification. Settings for DIA-NN are as follows: output filtered at 0.01 false-discovery rate (FDR); generate spectral library option selected (deep learning was used to generate a new in silico spectral library from peptides provided); library-free search enabled; minimum fragment *m/z* set to 200 and maximum fragment *m/z* set to 2000 (based on instrument MS2 settings); N-terminal methionine excision enabled; in silico digest cut at K*,R* but excluded cuts at *P (since only trypsin was used, and not a combination of trypsin with LysC); maximum number of missed cleavages set to 1; minimum peptide length set to 7; maximum peptide length set to 30; minimum precursor m/z set to 399 and maximum precursor *m/z* set to 1200 (based on DIA window scheme); minimum precursor charge set to 1; maximum precursor charge set to 4; Cysteine carbamidomethylation enabled as a fixed modification; a double pass mode neural network classifier was used were used for peak selection; the smart profiling library generation option was used such that a spectral library was also created from the DIA runs and used to reanalyze them. When generating a spectral library, *in silico* predicted spectra were retained if deemed more reliable than experimental ones; DIA-NN was set to optimize the mass accuracy automatically using the first run in the experiment. Since DIA-NN was made to handle protein inference and grouping assuming UniProtKB formatted FASTA, and the NCBI RefSeq *Desmodus rotundus* Release 100 was used (RefSeq not UniProtKB), settings were chosen such that DIA-NN effectively ignored protein grouping, which can be performed on the backend following ontology mapping. This was accomplished by setting implicit protein grouping to isoform IDs and the additional option of "--relaxed-prot-inf". Lastly, for match between runs (MBR) and quantification strategy, the Robust LC (high precision) setting was chosen as well as retention time-dependent cross-run normalization (these retention time windows are automatically set during training). Resulting spectral libraries, result tables and summary reports from DIA-NN are deposited to the ProteomeXchange Consortium via the PRIDE (Perez-Riverol et al., 2022) partner repository with the dataset identifier PXD031075.

The serum protein identifications were converted to human orthologs to aid in downstream analysis. This was accomplished by using a series of python scripts (Anaconda v2019.07; conda v4.7.11; Python v3.6.8) from GitHub on the pwilmart/PAW_BLAST and pwilmart/annotations repositories, retrieved November 18, 2019. Broadly, these tools take a list of identifiers to create a subset FASTA (make_subset_DB_from_list_3.py), which is then searched against a human FASTA (db_to_db_blaster.py; against the UniProtkKB Human canonical reference proteome UP000005640, release 2021_03) using a local installation of BLAST+ 2.11.0. (Camacho et al., 2009). The 652 experiment-wide bat protein identifications (i.e., from both the primary study and the heat inactivation study) were assigned human orthologs, which were then manually inspected for incorrect assignments. Such assignments can happen when a protein is not present in humans, or if the BLAST hit disagrees with the original bat annotation, in which case the original annotation is preserved (e.g., pregnancy zone protein is frequently incorrectly mapped to alpha-2-macroglobulin following BLAST). In cases where human orthologs do not exist, such as mannose-binding protein A (MBL1), we used ad hoc ortholog identifiers. Specifically, eight identified vampire bat proteins are not present in humans (MBL1, APOR, HBE2, REG1, LGB1, Bpifb9a, ICA, and Patr-A), and UniProt identifiers from chimpanzee, cow, horse, mouse, and pig were used instead. Finally, duplicate entries were summed together into a single non-redundant entry, and the final list of 633 mapped protein identifications, with links to entries on UniProtKB, are reported in Table S2.

Given prior proteomic identification of putative viral proteins in undepleted serum, including CoVs (Neely et al., 2020), we also broadened our search space for any CoV proteins. We performed a secondary search using the same settings and the addition of a Coronaviridae FASTA (117709 sequences) retrieved from UniProtKB (2021_03 release) using taxon identifier 1118 with all SwissProt and TrEMBL entries. Search settings and FASTA are included in the PRIDE submission (PXD031075). We identified seven CoV proteins at the 1% protein FDR level. These included 749 peptide identifications across all injections (including heating experiment and pool; Table S6). Given that observing non-host proteins is a rare event, we used additional stringent criteria to verify these identifications. We *a priori* defined thresholds of the following DIA-NN peptide-level scores: PEP < 0.01 (posterior error probability), MS1 Profile Correlation > 0.9, and Mass Evidence > 1. These are outputs are in Table S6 and detailed definitions are available at https://github.com/vdemichev/DiaNN#main-output-reference. Of the 749 CoV peptides, 483 had a PEP ≥ 0.01, and the remaining had MS1 Profile Correlation scores < 0.9. In this case, the criteria of Mass Evidence > 1 was never used. Overall, there were no viral proteins putatively identified in the undepleted serum, regardless of CoV infection status determined by PCR.

**S3 Supplemental tables**

Table S1. Data file key

Table S2. Experiment-wide protein identifications and relative abundance

Table S3. Heat inactivation experiment protein abundance and statistics

Table S4. CoV experiment protein abundance and statistics

Table S5. Serum protein ranks compared to Neely et al. 2020

Table S6. Extended metrics for CoV peptide spectral matches

**S3 Supplemental figures**

Figure S1. Overlap between vampire bat serum proteome composition from our 2020 analysis (sera collected in 2015) (Neely et al., 2020) and the current study (sera collected in 2019).

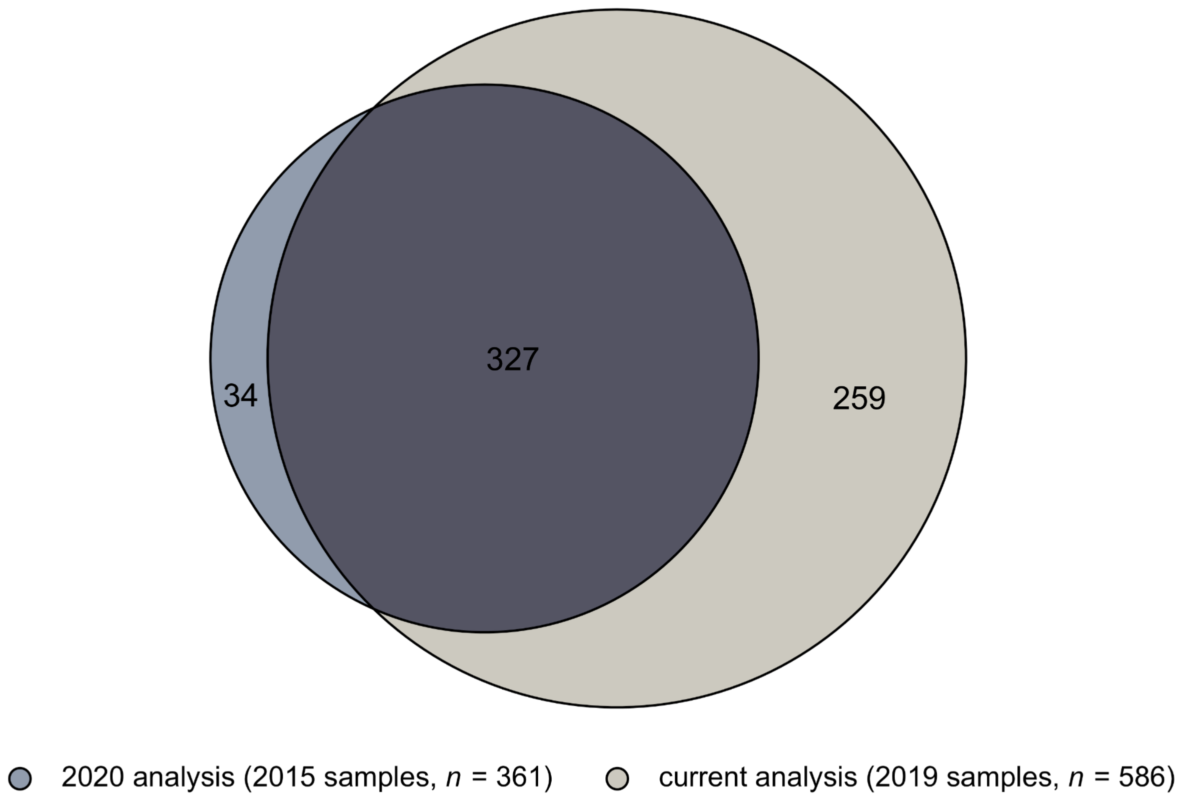

Figure S2. Distribution of mean absolute percent change for all proteins in response to heat inactivation among the four bats with paired pre- and post-treatment sera ($log_{10}$ scale). The black line shows the cumulative frequency (and the horizontal dashed line shows 50%). The vertical dashed line shows the mean absolute percent change including 50% of proteins (~16.3%).

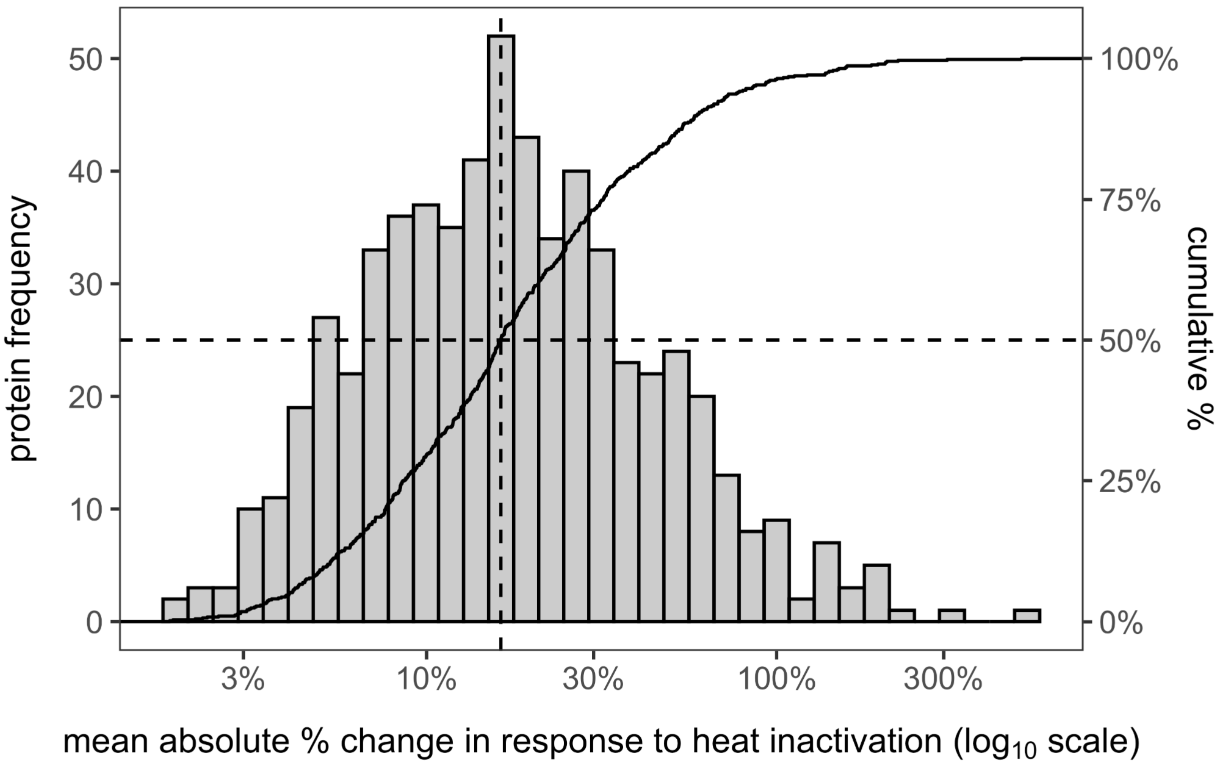

Figure S3. Biplot of the first two principal components (PCs) from the PCA of the 586 identified serum proteins. Missing abundance values were imputed as half the minimum intensity per protein. Individual vampire bats are colored by their CoV infection status, and ellipses display the standard error of CoV group centroids using the *ggordiplot* package.

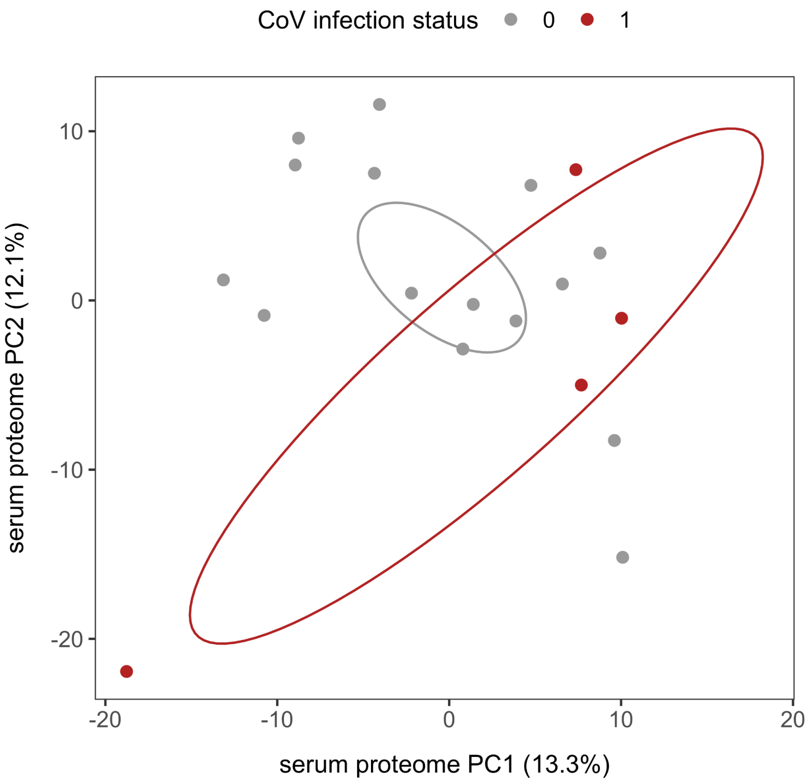

Figure S4. Top GO biological processes in the vampire bat serum proteome, defined as any process (accessed programmatically using the *UniprotR* package) with at least 30 proteins. Processes are ordered by their representation in the proteome (*n* = 586 identified proteins).

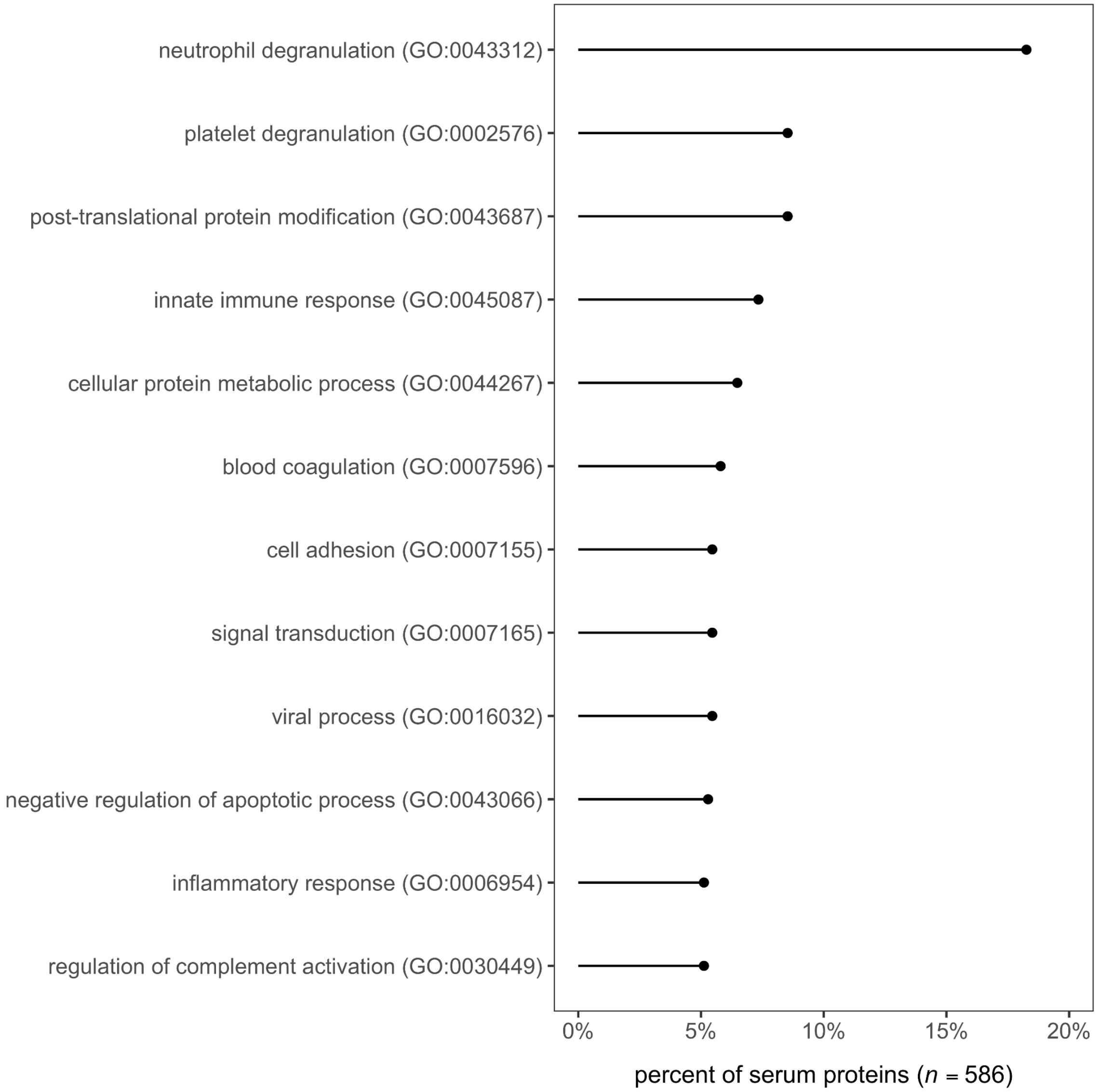
